# Sex-dependent metabolic remodeling of kidneys revealed by arteriovenous metabolomics

**DOI:** 10.1101/2024.09.02.610869

**Authors:** Miranda E. Kelly, Lauren A. Hoffner, Cuauhtemoc B. Ramirez, Alexis L. Anica, Joohwan Kim, Gregory Tong, Yeojin Kim, Wonsuk Choi, Kihong Jang, Yasmine H. Alam, Sunhee Jung, Johnny Le, Ian Tamburini, Miranda L. Lopez, Hosung Bae, Yujin Chun, Won-Suk Song, Thomas F. Martinez, Cholsoon Jang, Gina Lee

**Affiliations:** Department of Biological Chemistry, Center for Epigenetics and Metabolism, Chao Family Comprehensive Cancer Center, University of California Irvine School of Medicine, Irvine, CA 92697-4025, USA; Department of Microbiology and Molecular Genetics, Center for Epigenetics and Metabolism, Chao Family Comprehensive Cancer Center, University of California Irvine School of Medicine, Irvine, CA 92697-4025, USA; Department of Pharmaceutical Sciences, Chao Family Comprehensive Cancer Center, University of California Irvine School of Pharmacy and Pharmaceutical Sciences, Irvine, CA 92697-4025, USA

## Abstract

Sex is a fundamental biological variable important in biomedical research, drug development, clinical trials, and prevention approaches. Among many organs, kidneys are known to exhibit remarkable structural, histological, and pathological differences between sexes. However, whether and how kidneys display distinct metabolic activities between sexes is poorly understood. By developing kidney-specific arteriovenous (AV) metabolomics combined with transcriptomics, we report striking sex differences in both basal metabolic activities and adaptive metabolic remodeling of kidneys after a fat-enriched ketogenic diet (KD), a regimen known to mitigate kidney diseases and improve immunotherapy for renal cancer. At the basal state, female kidneys show highly accumulated aldosterone and various acylcarnitines. In response to the KD, aldosterone levels remain high selectively in females but the sex difference in acylcarnitines disappears. AV data revealed that, under KD, female kidneys avidly take up circulating fatty acids and release 3-hydroxybutyrate (3-HB) whereas male kidneys barely absorb fatty acids but consistently take up 3-HB. Although both male and female kidneys take up gluconeogenic substrates such as glycerol, glutamine and lactate, only female kidneys exhibit net glucose release. Kidney transcriptomics data incompletely predict these sex differences, suggesting post-transcriptional/translational regulation mechanisms. This study provides foundational insights into the sex-dependent and diet-elicited metabolic flexibility of the kidneys in vivo, serving as a unique resource for understanding variable disease prevalence and drug responses between male and female kidneys.

## Introduction

Sex is considered a critical factor in preclinical and clinical studies. In nephrology, it has long been recognized that males and females exhibit distinct kidney histochemical structures. For example, the Na^+^-Cl^-^ co-transporter (NCC) encoded by *SLC12A3* is expressed at much higher levels in females than males, resulting in a greater response to thiazides, an NCC inhibitor prescribed as an antihypertensive drug^1^. Also, sex-specific differences in renal structure, renal sex hormone receptors, and gene expression have been reported and may contribute to differences in kidney functions^2^. For instance, the cell sex of proximal tubular epithelial cells (PTECs) affects kidney physiology, with male PTECs showing greater oxidative stress and apoptosis compared to female PTECs^3^. Another example is cuboidal cell transformation of the outer layer of the glomerular capsule observed in 60% of males but only in 5% of females^4^.

Women remain disproportionately under-represented in numerous studies regarding kidney physiology and pathophysiology. Few studies have reported sex differences in renal metabolite handling, gene expression, disease vulnerability and drug responses^2,5^. In rats, females display higher expression of sodium-glucose cotransporter (SGLT) 1 and 2, indicating a greater reliance on glucose transporters and potential differences in renal glucose handling between sexes^6^. Relatedly, women generally experience more severe adverse side effects such as ketoacidosis when prescribed SGLT2 inhibitors to treat type II diabetes^7^.

While kidneys have long been recognized for their critical role in blood filtration and waste excretion, an equally important role of kidneys is to maintain circulating metabolite homeostasis by generating and releasing various metabolites into the bloodstream. For instance, along with the liver, kidneys are major contributors to blood glucose and ketone levels through gluconeogenesis and ketogenesis. Kidneys also control whole-body acid-base homeostasis. In addition, kidneys synthesize and release select amino acids such as arginine, serine and glycine^8^. Lastly, there are kidney-specific metabolic products such as guanidinoacetate, the precursor of creatine^9–11^. Thus, elucidating sex differences in renal metabolism will be instrumental in developing more effective strategies for treating kidney diseases.

In this regard, the fat-enriched ketogenic diet (KD) has gained recent interest due to its improving effects on kidney diseases and renal cell carcinoma^7,12–14^. However, how kidney metabolism changes upon KD and how such changes are distinct between sexes remain elusive. Thus, we sought to characterize sex-specific renal metabolic profiles under normal diet and KD conditions. Here, by developing kidney-specific AV metabolomics in mice, we present a comprehensive landscape of metabolite trafficking in male and female kidneys, revealing that biological sex significantly impacts renal metabolism both at the basal state and in response to KD. Altogether, these findings provide novel insights into the impact of sex, as a critical biological variable, on renal metabolism.

## Results and Discussion

### Sex differences in kidney metabolome

To determine sex-dependent renal metabolism under different dietary conditions, we began our study by comparing metabolomic differences between male and female kidney tissues from 10-week-old C57BL/6 mice fed either normal chow (NC) or ketogenic diet (KD) (**Fig. 1A**). After 5 weeks of KD feeding, both males and females exhibited a similar increase in the circulating levels of the ketone body 3-HB, reflecting increased fatty acid oxidation^13^ (**Fig. 1B**). Then, ad libitum fed mice were sacrificed in the early morning without fasting to maximize the diet effect, and metabolites were extracted from kidneys for liquid chromatography-mass spectrometry (LC-MS) analysis. From our metabolomics platform with annotated metabolites confirmed by authenticated chemical standards, we identified 232 significantly different metabolites between sexes and diets (**Fig. 1C**).

Volcano plots showing sex differences in renal metabolite abundances indicated that, under NC-fed conditions, 133 metabolites were higher in female kidneys while 61 metabolites were higher in male kidneys (**Fig. 1D**). Aldosterone, a hormone that regulates the kidney’s sodium and water reabsorption, was ∼18-fold higher in female kidneys than in male kidneys, regardless of diet (**Fig. 1E**). In addition, various acyl-carnitine species, products of fatty acid oxidation, showed female-specific renal accumulation (**Fig. 1F, G**), suggesting differential fatty acid oxidation activities between sexes. However, carnitine itself did not exhibit such sex-specific patterns (**Fig. 1H**).

The sex differences in kidney metabolome were substantially dampened by KD (**Fig. 1I**), with only 68 and 39 metabolites being distinctly abundant between sexes. While higher aldosterone levels were still observed in female kidneys under KD conditions (**Fig. 1E**), female-specific renal accumulation of acyl-carnitines was no longer noticed (**Fig. 1I, J**). Further, sex differences in metabolite abundances across multiple classes, including amino acids and organic acids, were also diminished by KD (**Fig. 1D, I**). Thus, KD has a major impact on renal metabolome regardless of sex.

### Sex difference in metabolite trafficking during diet switching

Since metabolic differences in the kidney between sexes were seen as early as 5 weeks on KD, we sought to track the progression of such sex-dependent metabolic rewiring during diet switching. To this end, we systematically compared 0, 3, and 5 weeks of KD-fed male and female kidneys. Circulating 3-HB measurements indicated that 3-week KD-fed mice can serve as an intermediate metabolic transition state between 0 and 5-week KD-fed mice in both sexes (**Fig. 2A**).

In addition to this multiple time-frame investigation, we implemented additional technical innovation to elucidate metabolite movements across the kidney, given that snapshot metabolite levels do not often provide metabolic flux information. We therefore developed a kidney-specific arterio-venous (AV) metabolomics technique to interrogate the impact of sex and diet on the kidney’s metabolite trafficking activities or fluxes (i.e., uptake and release of metabolites by the kidney). To this end, arterial (A) blood and renal venous (RV) blood were sampled and metabolite abundances between these two blood samples were compared to quantitate the kidney’s dynamic uptake (RV-A<0) and release (RV-A>0) of circulating metabolites (**Fig. 2B**). Our successful establishment of this technique in mice was validated by the detection of renal release of guanidinoacetate, a creatine precursor synthesized solely by the kidney^9,15^ (**Fig. 2C**), and the renal uptake of citrate, a fuel source preferentially taken up by the kidney^10^ (**Fig. 2D**). This kidney-specific activity was consistently observed across different sexes and diets.

**Figure 1.**
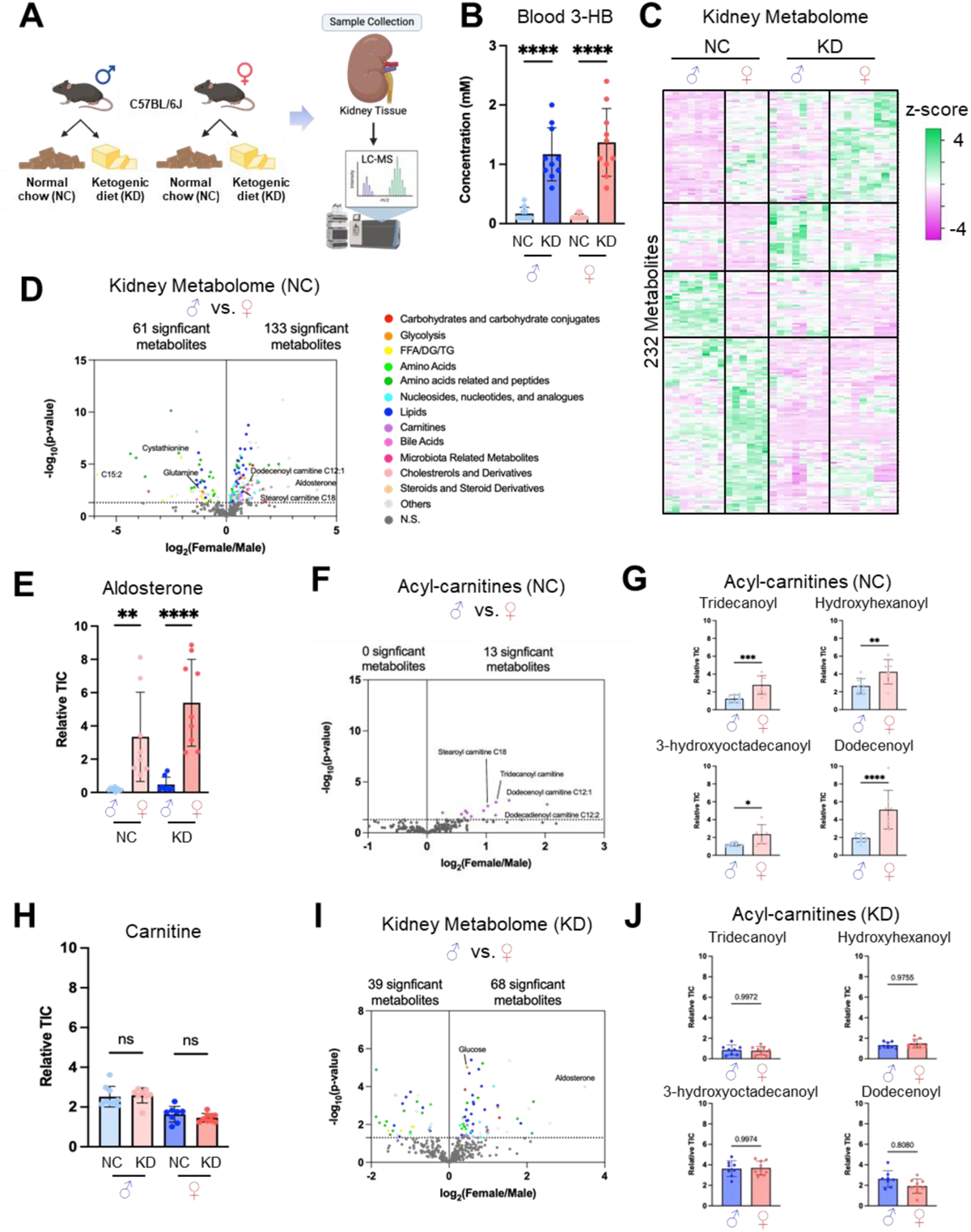
Sex differences in kidney metabolome. (A) Experimental scheme of male and female mice after NC or KD feeding for 5 weeks. (B) Blood 3-HB levels. (C) Heatmap showing 232 kidney metabolites different between sexes and diets. (D) Volcano plot showing kidney metabolites different between sexes in NC. Colors indicate various categories of metabolites. (E) Kidney aldosterone abundance. (F) Volcano plot showing kidney acyl-carnitines different between sexes in NC. (G) Kidney acyl-carnitine abundance in NC. (H) Kidney carnitine abundance. (I) Volcano plot showing kidney metabolites different between sexes in KD. (J) Kidney acyl-carnitine abundance in KD. N=8 and N=9 mice for ND-fed and KD-fed females. N=9 and N=8 mice for ND-fed and KD-fed males. *p<0.05, **p<0.01, ***p<0.001, ****p<0.0001 by Students’ t-test. n.s.; not significant.

The AV metabolomics analysis revealed that, in both sexes and during diet switching, the kidney took up a higher number of metabolites than it released (**Fig. 2E-J**). Among them, the most consistently observed kidney activity was the uptake of TCA cycle intermediates (citrate, succinate, malate, fumarate and oxoglutarate) (**Fig. 2D and S1A**), suggesting their role in fueling the kidney’s energy metabolism and biosynthesis, regardless of sex and diet. In addition, the AV data further disclosed consistent renal uptake of previously unrecognized metabolites such as glutathione and hypotaurine in both sexes and diets (**Fig. S1B**). These sulfate-containing metabolites exert antioxidant activities, suggesting their utilization by the mitochondria-rich kidney to ameliorate oxidative stress from active oxidative phosphorylation^16^. In terms of metabolites released by the kidney, the most prominent ones were nucleotides (e.g., thymidine) and amino acids (e.g., glutamate, cystine) (**Fig. S1C**).

**Figure 2.**
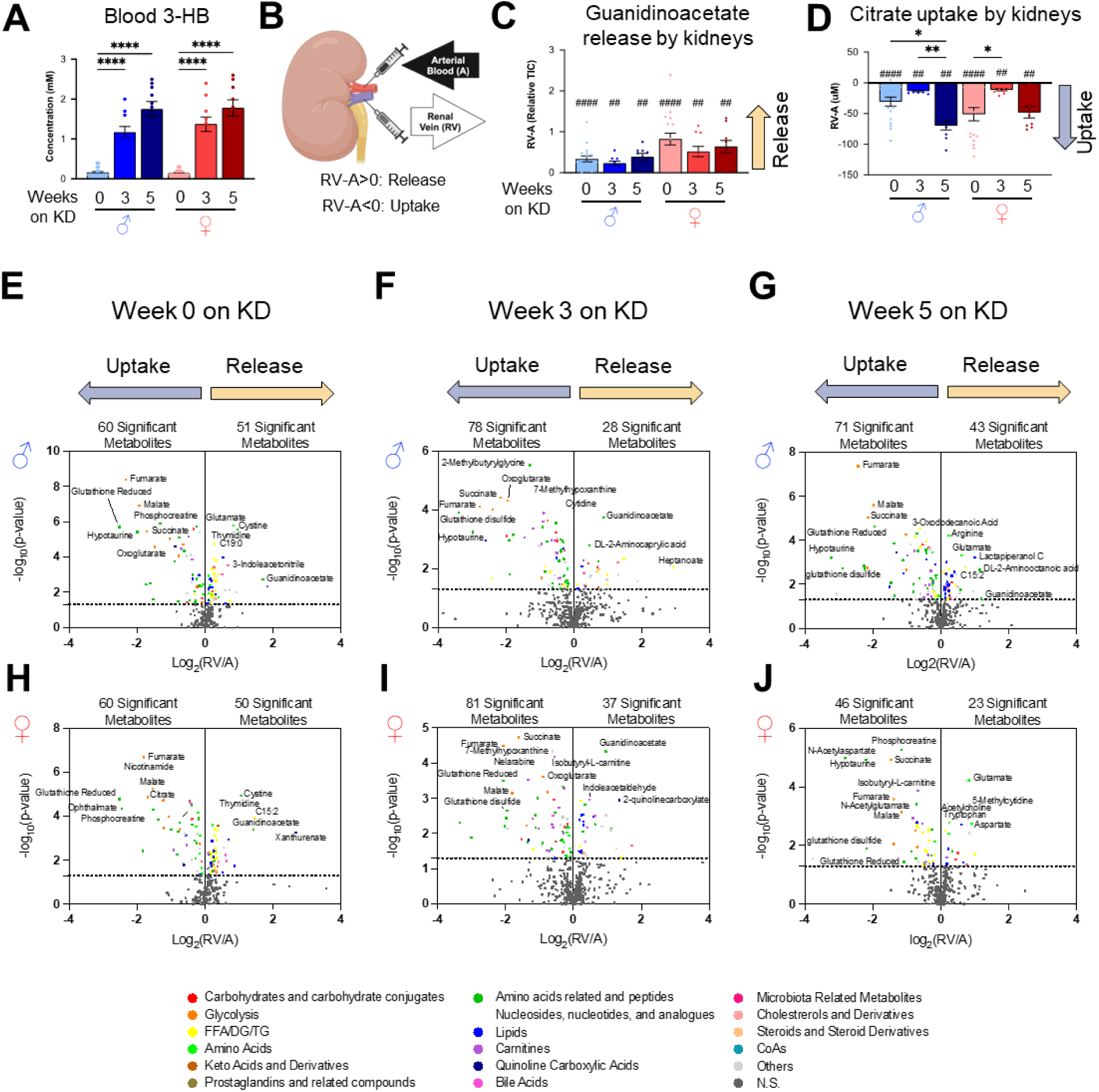
Sex differencse in metabolite trafficking during diet switching. (A) Blood 3-HB levels at weeks 0, 3, and 5 after diet switching. (B) Schematic of kidney-specific AV metabolite gradient analysis. (C, D) Consistent kidney release of guanidinoacetate (C) or uptake of citrate (D) across sexes and diets. (E-J) Volcano plots showing metabolites whose kidney uptake and release are significantly different in males (E-G) and females (H-J) or during diet switching from week 0 (E, H), week 3 (F, I) and week 5 (G, J). N=18, 9, and 9 mice for 0, 3 and 5 weeks KD-fed males and N=17, 10, and 9 mice for 0, 3, and 5 weeks KD-fed females. *p<0.05, **p<0.01, ****p<0.0001 by Students’ t-test. ^#^p<0.05, ^##^p<0.01, ^####^p<0.0001 by one-sample t-test.

**Figure 3.**
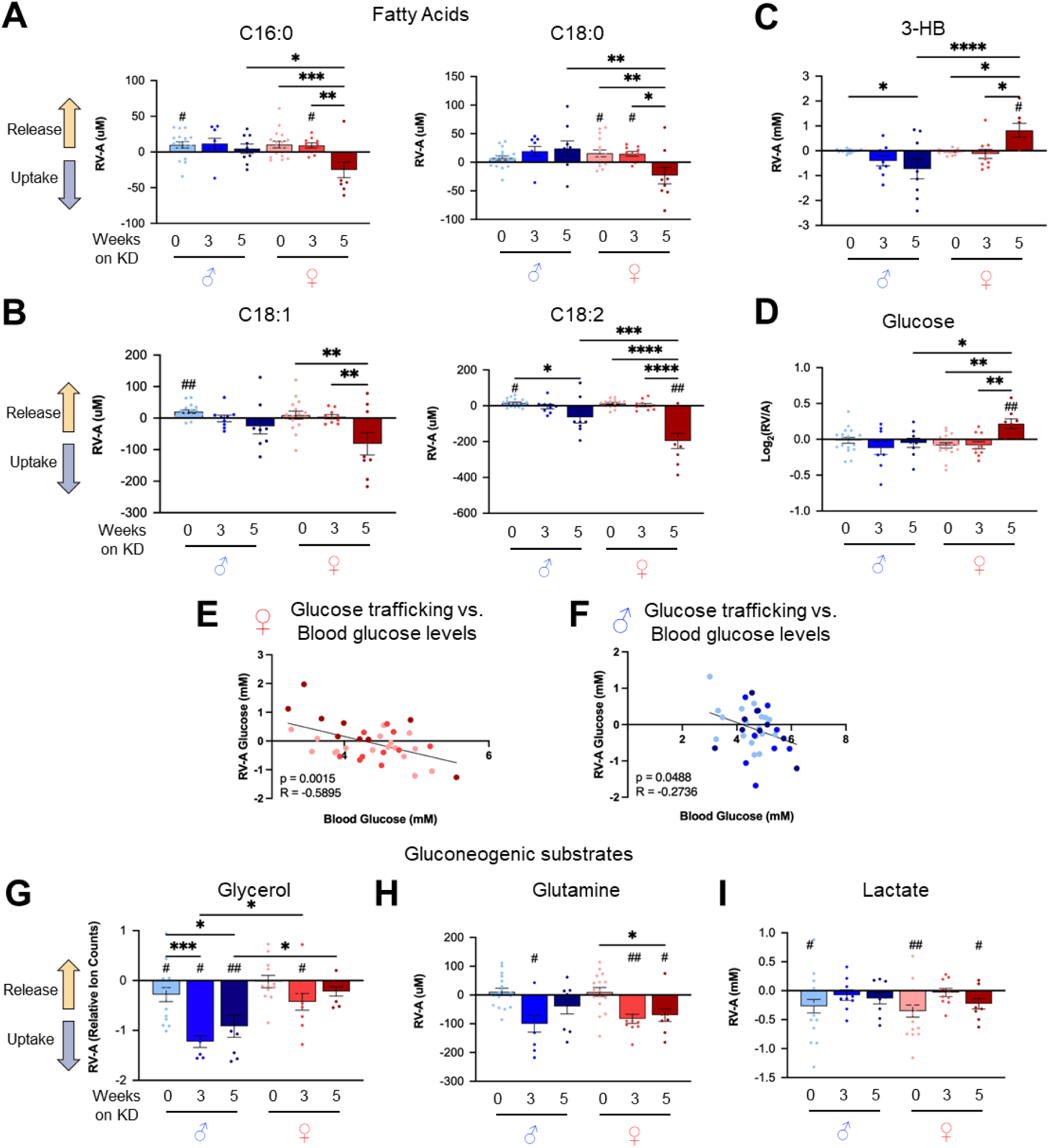
Sex-dependent ketogenesis and gluconeogenesis activities of the kidney. (A, B) Kidney release and uptake of saturated (A) and unsaturated (B) free fatty acids. (C, D) Kidney release and uptake of 3-HB (C) and glucose (D). (E, F) Correlation between blood glucose concentrations and kidney glucose release or uptake in females (E) or males (F). (G-I) Kidney release and uptake of gluconeogenic substrates. N=18, 9, and 9 mice for 0, 3 and 5 weeks KD-fed males and N=17, 10, and 9 mice for 0, 3, and 5 weeks KD-fed females. *p<0.05, **p<0.01, ***p<0.001, ****p<0.0001 by Students’ t-test. ^#^p<0.05, ^##^p<0.01 by one-sample t-test. Pearson correlation test was used in E and F.

Notably, upon diet switching to KD, we observed progressive changes in the trafficking of fatty acids and lipid species across kidneys (from release toward uptake after diet switching, highlighted yellow and dark blue in the volcano plots) (**Fig. 2E-J**). Nevertheless, the kind of metabolites and the extent of trafficking were markedly distinct between sexes and dynamically changed during diet switching (see below).

### Sex-dependent ketogenesis and gluconeogenesis activities of the kidney

Based on the observed sex differences in fatty acids, lipids, and acyl-carnitine abundances and trafficking, we first sought to determine the trafficking patterns of the 4 most abundant circulating fatty acids that not only serve as fuels but also building blocks for lipids and acyl-carnitines. Upon diet switching from 0, 3 to 5 weeks, male kidneys showed no significant changes in the trafficking of saturated fatty acids, palmitate (C16:0) and stearate (C18:0) (**Fig. 3A**). In contrast, they gradually increased uptake of unsaturated fatty acids, oleate (C18:1) and linoleate (C18:2) (**Fig. 3B**). On the other hand, female kidneys potently absorbed all four fatty acids after 5 weeks of KD but not after 3 weeks (**Fig. 3A, B**), suggesting delayed metabolic remodeling of the kidney in females.

Fatty acids are the major substrates of ketogenesis, and alongside the liver, the kidney is the primary organ responsible for synthesizing and releasing ketone bodies into circulation^17–19^. We thus examined the trafficking of 3-HB and found striking sex differences: males increased the uptake of 3-HB gradually over time during diet switching, while females released 3-HB only after 5 weeks of KD (**Fig. 3C**). This time-frame was reminiscent of unsaturated fatty acid trafficking (**Fig. 3B**), suggesting the link between these two metabolic processes.

In addition to ketogenesis, the kidney is known to have a gluconeogenesis capacity, contributing to whole-body glucose homeostasis, particularly after starvation or in individuals with diabetes^18,20–22^. We thus examined kidney gluconeogenesis during the diet switching and observed that glucose release was prominent only in female kidneys upon the KD feeding (**Fig. 3D**). However, no significant glucose release was observed in male kidneys (**Fig. 3D**). Notably, renal glucose release by female kidneys was negatively correlated with blood glucose concentrations, with more glucose being released when blood glucose levels were lower (**Fig. 3E**). This finding suggests homeostatic mechanisms by which the body senses circulating glucose levels and controls kidney’s glucose release to maintain systemic glucose concentrations^22–24^. A similar but weaker relationship was observed in males (**Fig. 3F**).

Next, we were curious whether the observed sex difference in gluconeogenesis originated from different availability of gluconeogenic substrates. We thus measured the renal uptake of three key gluconeogenic substrates in circulation: glycerol, glutamine and lactate (**Fig. 3G-I**). For glycerol measurement, we employed a chemical derivatization method using glycerol kinase followed by LC-MS analysis of glycerol-3-phosphate because glycerol is not measurable by typical LC-MS^25,26^. Unexpectedly, male kidneys showed more glycerol uptake than female kidneys (**Fig. 3G**) despite their absence of glucose release (**Fig. 3D**). In addition, male kidneys but not female kidneys increased their uptake of glycerol upon KD, suggesting the alternative usage of glycerol, perhaps for lipid synthesis or oxidation rather than gluconeogenesis. In contrast, both male and female kidneys increased their uptake of glutamine to a similar extent 3 weeks after diet switching (**Fig. 3H**). On the other hand, lactate exhibited a trend of decreased uptake after diet switching in both sexes (**Fig. 3I**), likely reflecting overall decreased glucose utilization and lactate production upon KD. Thus, we concluded that distinct gluconeogenesis activity between male and female kidneys is not due to the availability of circulating gluconeogenic substrates, but rather is associated with sex-dependent kidney metabolism per se.

**Figure 4.**
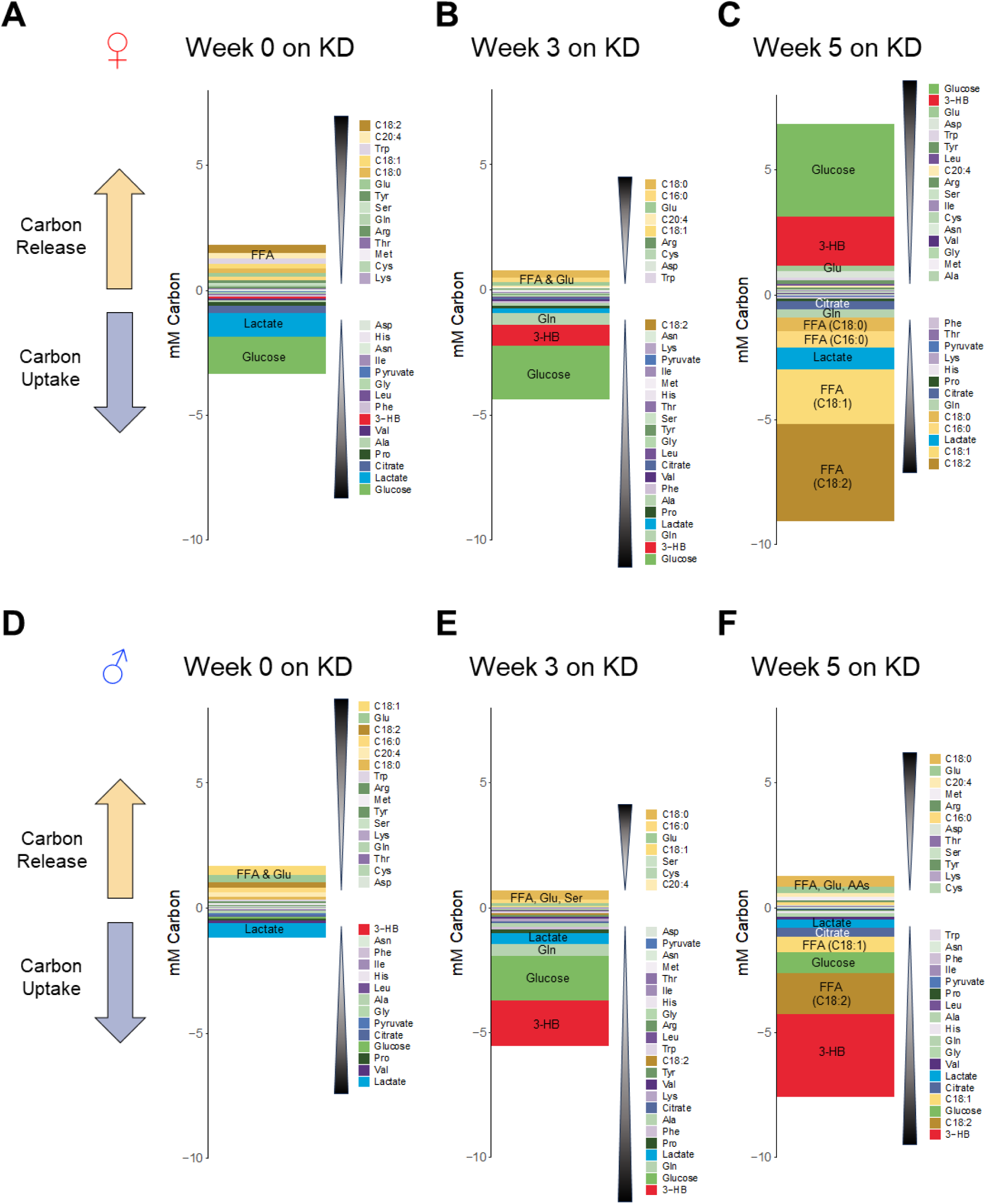
Quantitative analysis of kidney carbon trafficking. (A-F) Carbon release and uptake via each metabolite by female kidneys (A-C) and male kidneys (D-F) during diet switching on week 0 (A, D), week 3 (B, E) and week 5 (C, F). Metabolites are ordered based on their relative contributions from greatest to least.

### Quantitative analysis of kidney carbon trafficking

Carbon is the most fundamental element for living cells, yet the quantitative understanding of carbon usage by kidneys is limited. Using our AV data, we thus performed a quantitative analysis of total carbon uptake and release by kidneys in both sexes and during diet switching. To this end, we determined the absolute concentrations of 30 major circulating nutrients using standard curves generated with authentic chemical standards and calculated the concentration gradients of total carbons carried by each metabolite based on its chemical formula (e.g., six carbons for glucose, sixteen carbons for palmitate, see methods for detail). In female kidneys prior to diet switching (week 0), the primary carbon carriers were circulating glucose and lactate, accounting for ∼45% and 35% of total carbon influx (**Fig. 4A**). Remarkably, citrate also contributed a significant portion (∼10%) of the total carbon influx. On the other hand, several fatty acids and amino acids carried the most carbon efflux, with each accounting for ∼50%. In this condition, the total carbon influx and efflux were well balanced. On week 3 after diet switching, kidneys still took up a large amount of carbons as glucose (∼50%) but much less as lactate. Instead, 3-HB (∼25%) and glutamine (∼15%) became significant contributors to total carbon influx (**Fig. 4B**). In this condition, we observed a decrease in carbon release as fatty acids and amino acids, resulting in higher carbon influx than efflux.

The most striking change in female kidneys occurred on week 5 after diet switching (**Fig. 4C**). The kidneys exhibited a profound increase in carbon uptake from fatty acids, leading to ∼3-fold increase in total carbon influx compared to week 0 (∼9 mM on week 5 versus ∼3 mM on week 0). Most of these carbons appeared to be converted to glucose and 3-HB, with an overall balanced total carbon influx and efflux. These analysis highlight substantial metabolic rewiring of female kidneys to augment gluconeogenesis and ketogenesis during diet switching.

In male kidneys, a different pattern emerged. In general, male kidneys on week 0 showed slightly less total carbon influx and efflux compared to female kidneys (**Fig. 4D**). Intriguingly, on week 3 after diet switching, male kidneys potently increased carbon influx through circulating glucose, 3-HB, and glutamine uptake (**Fig. 4E**). Despite an increase in carbon influx, carbon efflux did not increase. Accordingly, the balance of carbon influx and efflux was not achieved, similar to female kidneys on week 3 (**Fig. 4B**). These data suggest increased carbon storage within kidneys (as glycogen or lipids). Alternatively, there may be the release of other unmeasured carbon carriers such as lipids, proteins or CO_2_. On week 5, noticeable changes were the decreased carbon uptake as glucose and increased carbon uptake as 3-HB and unsaturated fatty acids (C18:1 and C18:2) (**Fig. 4F**). Together, these data provided quantitative insights into the contribution of circulating metabolites to kidney’s total carbon influx and efflux, which are altered during the diet switching and distinct between sexes.

### Quantitative analysis of kidney nitrogen trafficking

Next, we examined the kidney’s nitrogen handling, a fundamental yet less appreciated aspect of kidney metabolism. In female kidneys at week 0, nitrogen was taken up as both non-essential amino acids such as proline, glycine, and alanine and essential amino acids, mostly branched-chain amino acids (**Fig. 5A**). In contrast, the kidney released nitrogen mostly as non-essential amino acids, including arginine, glutamine, glutamate and serine. This pattern of nitrogen release, predominantly as non-essential amino acids, is logical because the loss of essential amino acids involves a catabolic process such as protein breakdown. In this regard, the release of nitrogen as tryptophan, an essential amino acid, was unique and unforeseen. Altered metabolism of tryptophan in various kidney diseases and the effect of kidney function on circulating tryptophan levels have been reported^27,28^. Thus, it will be important to elucidate the biological significance of tryptophan release by kidneys.

On week 3 after diet switching, female kidneys increased nitrogen uptake as glutamine while decreasing nitrogen release (**Fig. 5B**). This created an imbalance between nitrogen influx and efflux, which raised the possibility of nitrogen release through other metabolites such as ammonia or intact proteins. Kidneys serve as important endocrine organs by producing and releasing many protein hormones including erythropoietin, renin, kinins and gastrin^29,30^. Thus, kidneys may release hormones to help metabolic adaptations through inter-organ communications during diet switching. Kidneys may also utilize a significant amount of nitrogen for new protein synthesis to facilitate their own metabolic remodeling triggered by dietary switching. On week 5 after diet switching, the nitrogen balance was restored, driven by an increase in nitrogen release, primarily as non-essential amino acids, including aspartate, arginine and glutamate (**Fig. 5C**). This change may implicate the completion of the remodeling process, a notion further supported by the increased fatty acid uptake, ketone release and glucose release in female kidneys on week 5, but not on week 3 after diet switching (**Fig. 3A-D**).

The landscape of nitrogen trafficking in male kidneys was somewhat similar to female kidneys (**Fig. 5D-F**), contrasting with the distinct sex differences observed in carbon trafficking (**Fig. 4**). For example, male kidneys also increased nitrogen uptake as glutamine and decreased nitrogen release on week 3 (**Fig. 5D, E**). Again, this created a nitrogen imbalance between influx and efflux. However, one notable difference was the release of nitrogen as methionine by male kidneys on week 5 (**Fig. 5F**), which was not prominent in female kidneys (**Fig. 5C**). Methionine can be derived from protein breakdown, or de novo synthesized from homocysteine via betaine homocysteine methyltransferase (BHMT), an enzyme only expressed in the kidney cortex and liver^31,32^. Our RNA-seq analysis on the whole kidneys revealed sex differences in BHMT gene expression, with BHMT expressed exclusively in females while BHMT2 was significantly higher in males than females (**Fig. S2A, B**). This sex difference was conserved under KD (**Fig. S2A, B**). Thus, post-transcriptional mechanisms and/or other pathway genes are likely involved in male-specific methionine release.

**Figure 5.**
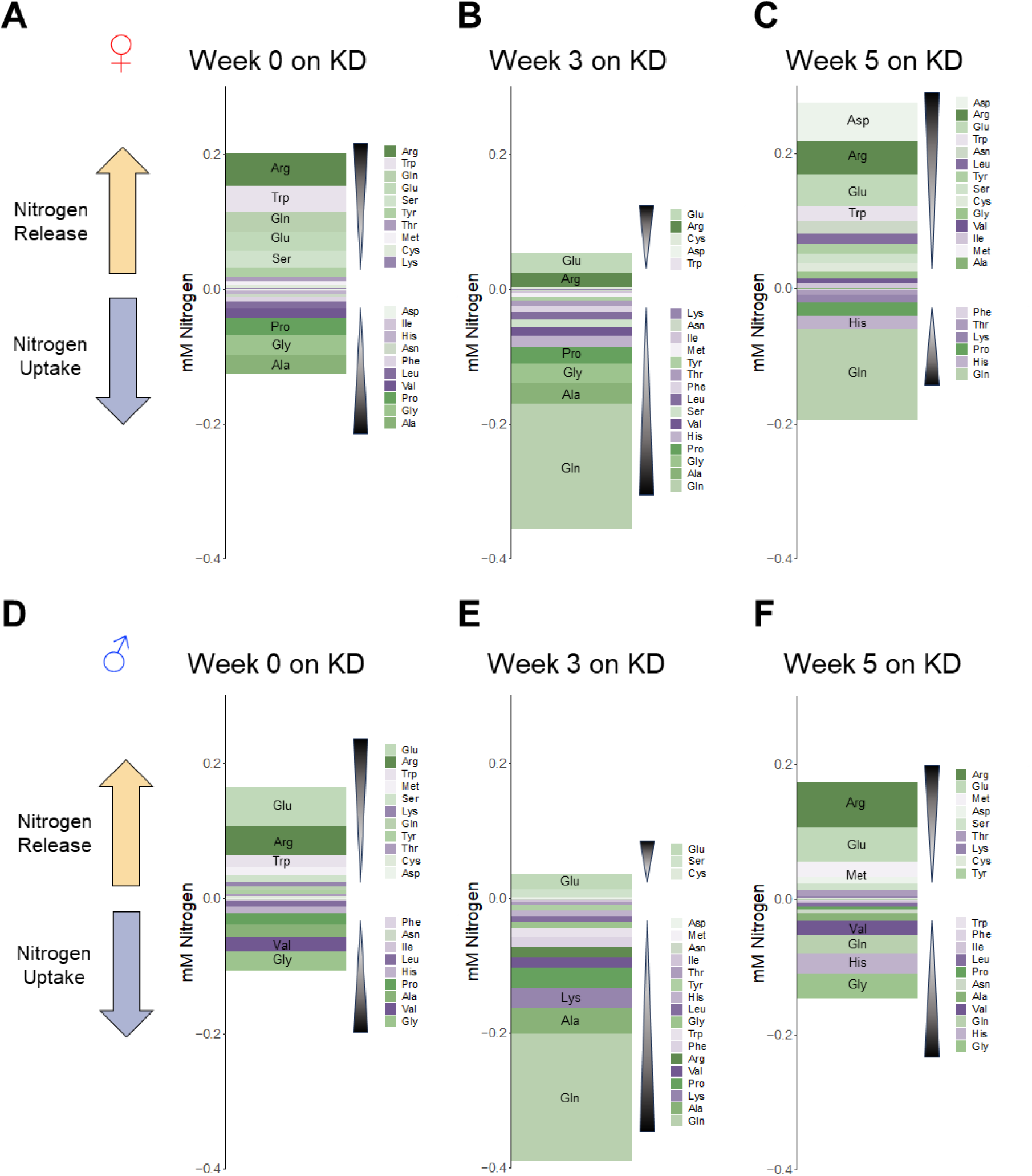
Quantitative analysis of kidney nitrogen trafficking. (A-F) Nitrogen release and uptake via each metabolite by female kidneys (A-C) and male kidneys (D-F) during diet switching on week 0 (A, D), week 3 (B, E) and week 5 (C, F). Metabolites are ordered based on their relative contributions from greatest to least.

### Sex-dependent kidney metabolic activities are largely unpredictable by gene expression

Finally, to determine whether sex-specific and diet-dependent metabolic activities of kidneys are transcriptionally regulated, we performed RNA-seq analysis of kidneys in both sexes and diets. The resulting data revealed that numerous genes were differentially expressed between sexes (**Fig. 6A**). Notably, we barely observed genes altered by KD in either sex (**Fig. 6A**). Principal component analysis further confirmed this notion (**Fig. S3A**). Thus, we decided to focused on genes influencing metabolic pathway activities (e.g., rate-limiting enzymes and metabolite transporters) to explain the observed sex differences and diet effects on renal metabolism.

In terms of ketogenesis, the rate-limiting enzyme is hydroxymethylglutaryl-CoA synthase 2 (HMGCS2) which converts acetoacetyl-CoA to 3-hydroxy-3-methylglutaryl-CoA (HMG-CoA), a precursor of 3-HB. Interestingly, on week 0, *Hmgcs2* gene expression was significantly higher in female kidneys than male kidneys and it was profoundly induced by KD in both sexes (**Fig. 6B**). Similarly, on week 5, *Hmgcs2* was significantly higher in female kidneys than male kidneys (**Fig. 6B**). However, this gene expression pattern appeared to be insufficient to explain the female kidneys’ 3-HB release and the male kidney’s 3-HB uptake (**Fig. 3C**).

Given that 3-HB trafficking was closely associated with fatty acid trafficking (**Fig. 3A-C**), we next surveyed genes for fatty acid uptake and oxidation that provide acetyl-CoA for ketogenesis. *Cd36*, a fatty acid transporter, showed higher expression in males than females, whereas Slc27a family fatty acid transporters (*Fatp1 and Fatp4*) were more highly expressed in females than males (**Fig. 6C**). However, KD did not affect their expression levels (**Fig. 6C**), raising the possibility that these transporters are not important drivers of ketogenesis in female kidneys and/or they are regulated by post-transcriptional mechanisms (e.g., membrane translocation). Carnitine palmitoyltransferases (Cpt1a and Cpt2), rate-limiting enzymes for fatty acid oxidation, also showed higher expression in males than females (**Fig. 6D**), which was not consistent with more active ketogenesis in female kidneys than male kidneys. Several long-chain acyl-CoA synthetases (ACSLs) that convert long-chain fatty acids to acyl-CoAs were expressed more highly in females than males but KD did not affect their expression (**Fig. 6E**). Collectively, these gene expression data did not consistently suggest more active ketogenesis in female kidneys than male kidneys.

Next, we examined the gene expression of gluconeogenic enzymes and transporters of gluconeogenic substrates. Phosphoenolpyruvate carboxykinase (PCK2) was the only gluconeogenic enzyme displaying higher gene expression in females although KD did not increase its levels (**Fig. 6F**). Other rate-limiting gluconeogenic enzymes exhibited either no sex difference (**Fig. S3B**) or male-enriched expression, for example, pyruvate carboxylate (PCX) (**Fig. 6F**). In terms of transporters for gluconeogenic substrates, only two among many monocarboxylate transporters, *Mct2* and *Mct4*, which import/export lactate and pyruvate, exhibited higher expression levels in female kidneys (**Fig. 6G**). Glutamine transporters also displayed higher expression levels in female kidneys (**Fig. S3C**), while glycerol transporters did not show sex differences (**Fig. S3D**). None of these transporters exhibited expression changes under KD. Thus, we concluded that post-transcriptional mechanisms play critical roles in sex differences in renal gluconeogenesis under KD.

**Figure 6.**
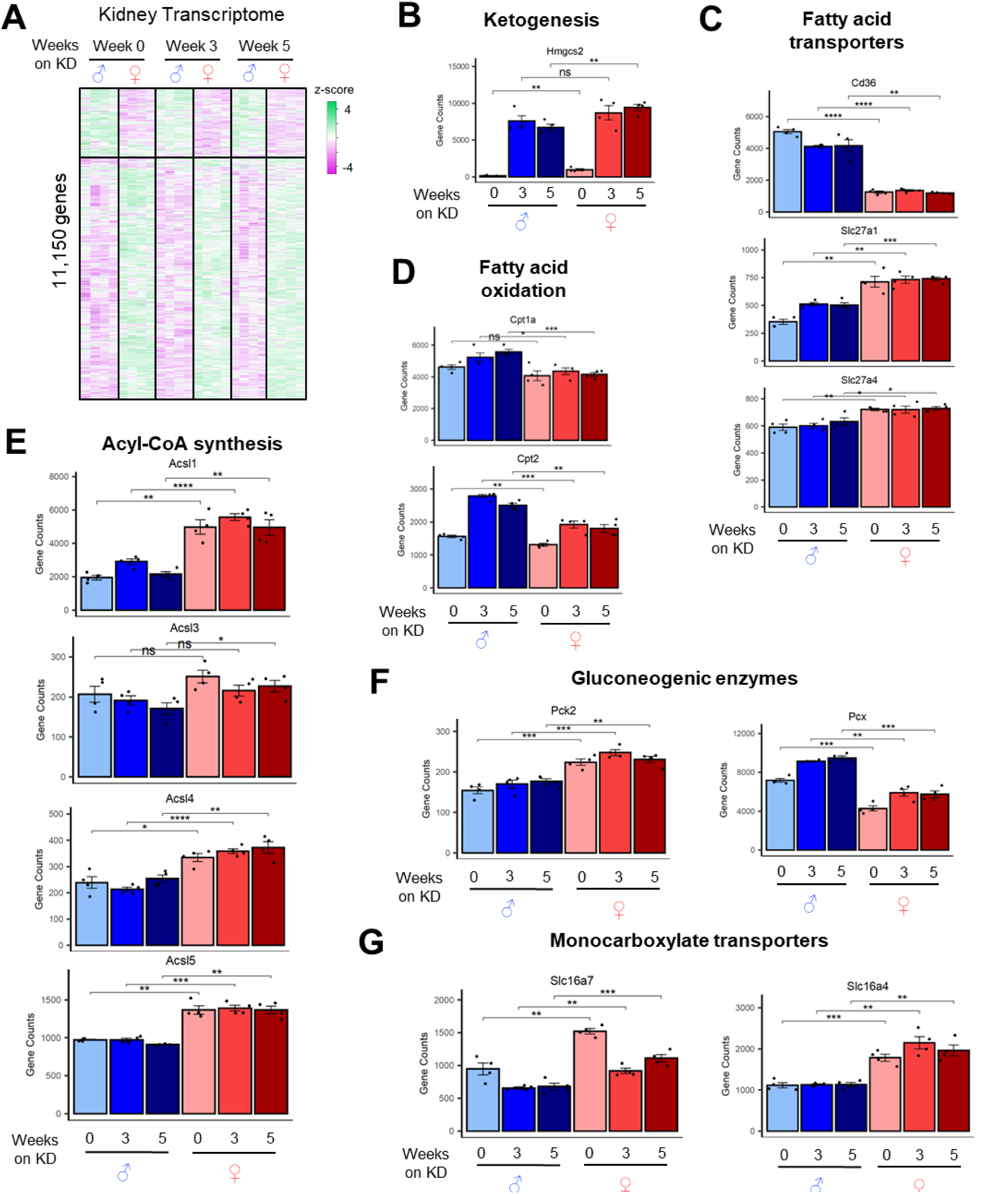
Sex-dependent kidney metabolic activities are largely unpredictable by gene expression. (A) Heatmap showing 11,150 kidney genes different between sexes and diets. N=4 per group. (B-G) Relative gene expression levels of the indicated enzymes and metabolite transporters relevant to ketogenesis and gluconeogenesis. N=4 per group. *p<0.05, **p<0.01, ***p<0.001, ****p<0.0001 by one-way ANOVA.

### Summary

In this study, we quantitate, for the first time, the vibrant metabolite trafficking activities of kidneys in both sexes during diet switching to KD. To achieve this, we developed a kidney-specific AV blood sampling technique followed by comprehensive metabolomics analysis. Our data uncovered markedly disparate basal kidney metabolism and responses to diet switching between males and females, especially female-specific ketogenesis and gluconeogenesis. Decreased renal ketogenesis and gluconeogenesis are well-known manifestations in patients with chronic kidney disease^33,34^. Thus, it will be important to determine the role of sexual dimorphism in renal ketogenesis and gluconeogenesis in these patient populations. Through comprehensive AV metabolomics, this study also reveals several metabolites previously unrecognized as being secreted or taken up by kidneys such as glutathione, hypotaurine and other poorly understood circulating metabolites, some of which likely have important signaling functions yet to be discovered. While we focused on ad-lib-fed conditions, fasting and feeding can influence kidney metabolic activities, making it crucial to perform AV metabolomics under these conditions. Kidneys have been reported to express lipoprotein lipase^35,36^. While our RNA-seq data indicated a strong reduction of lipoprotein lipase expression upon KD, kidneys may utilize fatty acids from lipoprotein-embedded triglyceride pools. AV lipidomics analysis will be able to quantitate kidneys’ usage of lipoproteins-derived lipids. Blood flow rate can impact absolute metabolic flux, although measuring blood flow to the kidney is technically challenging. How blood inputs to the kidney affect renal metabolism in a sex-dependent manner remains to be answered. In conclusion, this quantitative and systematic metabolic analysis offers new insights into distinct sex-and diet-dependent metabolic activities of kidneys, which will eventually help to understand the pathophysiological and clinical differences between males and females.

## Supplementary Figure Legends

**Figure S1.**
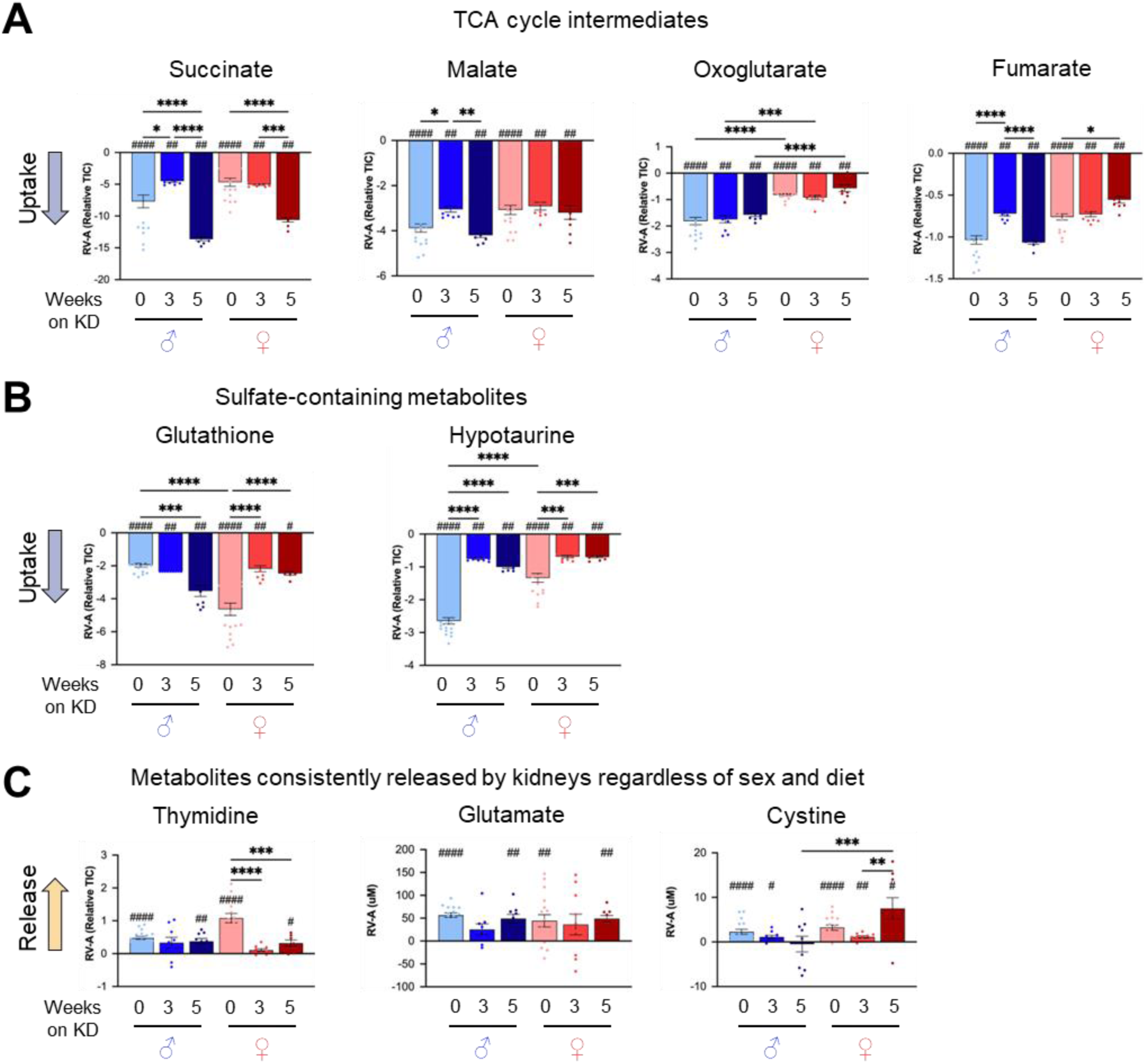
Metabolite trafficking during diet switching in male and female kidneys. (A-C) Kidney release and uptake of the indicated circulating metabolites. N=18, 9, and 9 mice for 0, 3 and 5 weeks KD-fed males and N=17, 10, and 9 mice for 0, 3, and 5 weeks KD-fed females. *p<0.05, **p<0.01, ***p<0.001, ****p<0.0001 by Students’ t-test. ^#^p<0.05, ^##^p<0.01, ^####^p<0.0001 by one-sample t-test.

**Figure S2.**
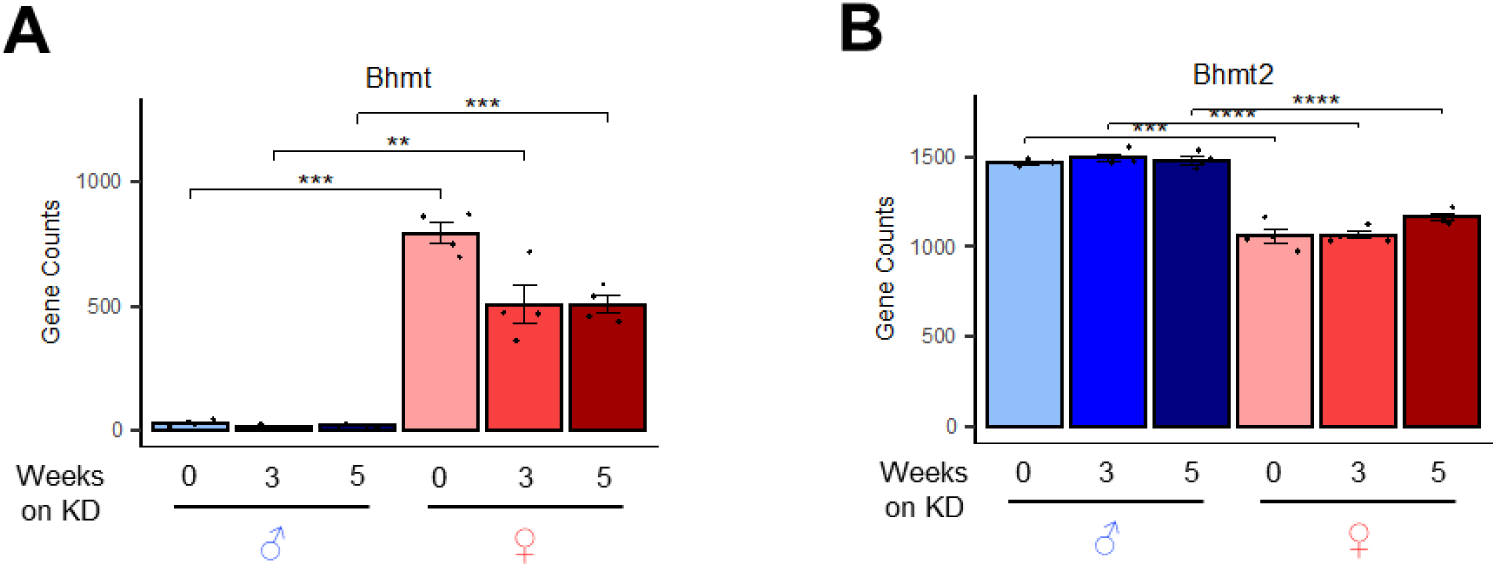
Sex differences in gene expression of *Bhmt* and *Bhmt2.* (A, B) RNA-seq gene counts of *Bhmt* or *Bhmt2* in male and female kidneys under NC or KD. N=4 mice per group. **p<0.01, ***p<0.001, ****p<0.0001 by one-way ANOVA.

**Figure S3.**
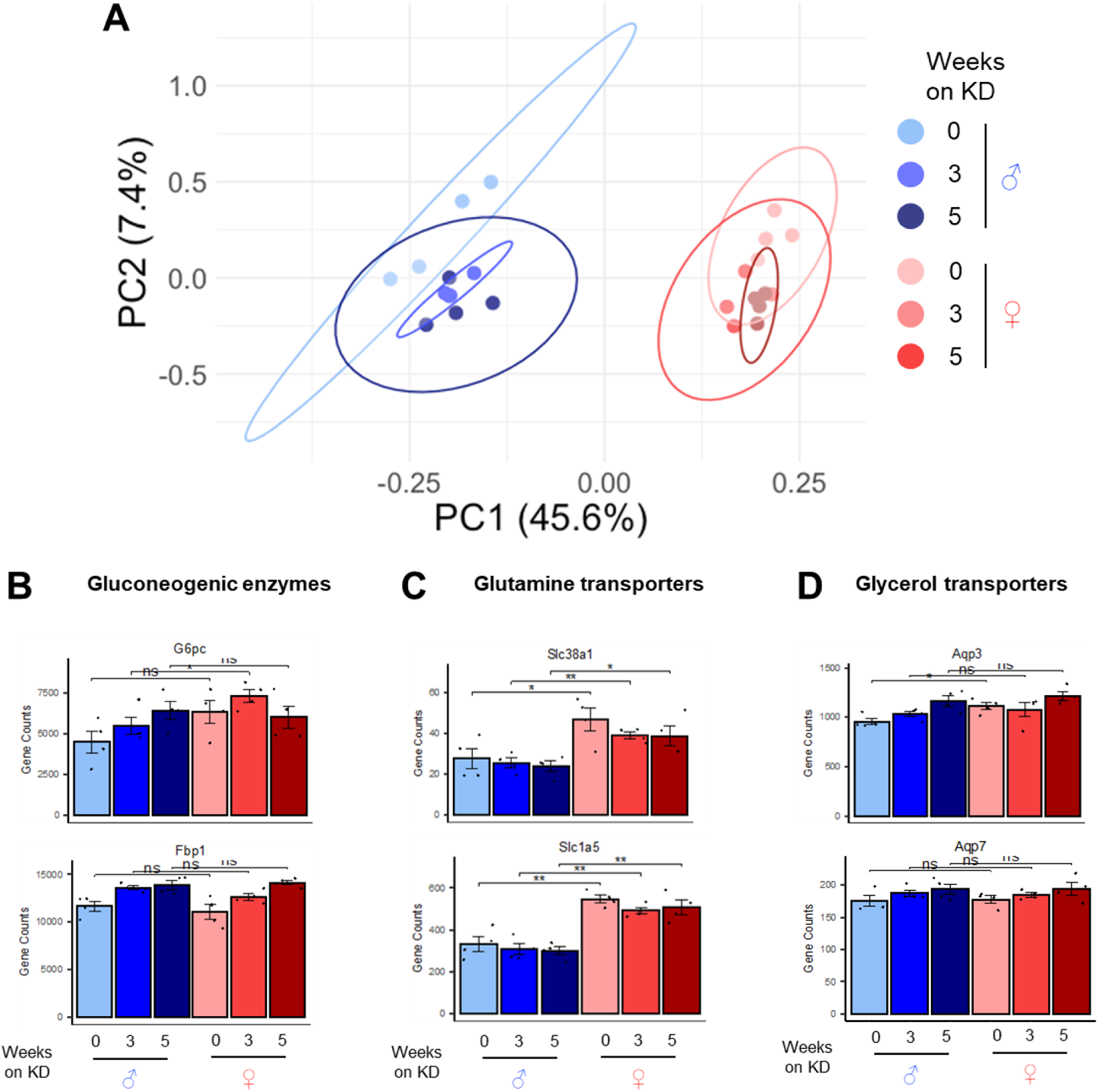
Sex differences in kidney transcriptome under NC and KD. (A) Principal component analysis (PCA) of RNA-seq results. (B-D) RNA-seq gene counts of the indicated genes. N=4 mice per group. *p<0.05, **p<0.01, by one-way ANOVA.

## Methods

### Mice

C57BL/6J male and female mice were purchased from Jackson Laboratories. Mice were housed at 22°C with a 12h light/12h dark cycle, free water access, and a chow diet. All animal experiments were approved by the University of California Irvine Institutional Animal Care and Use Committee. Mice were fed either normal chow or a ketogenic diet (Envigo, TD.160153) where carbohydrates were replaced with 44% Crisco, 15% cocoa butter, and 8% corn oil.

### Kidney AV blood sampling and tissue processing

At 8 AM, ad-lib-fed mice were anesthetized with isoflurane using the drop method and placed on a dissection tray with a nose cone. Blood collection was completed within 5 minutes post-anesthesia. The abdominal cavity was opened with a small incision and all abdominal organs were moved with fine surgical scissors to reveal the left kidney. The renal vein was identified and then punctured with an insulin syringe. ∼20 μL of blood was slowly collected with the syringe and transferred to an Eppendorf tube on ice, then the renal vein was clamped as the syringe was removed. Immediately after, the thoracic cavity was quickly opened by cutting through the ribcage and the aorta was cut with fine surgical scissors. ∼50 μL of pooled systemic blood in the thoracic cavity was collected with a cut pipet tip, transferred to an Eppendorf tube on ice. Blood samples were incubated on ice for 20 min, followed by centrifugation at 10,000 x g for 10 minutes at 4°C to obtain serum. The resulting supernatant was stored at - 80°C and analyzed within a week. The kidney tissues were snap-frozen using a liquid nitrogen-cooled Wollenburg clamp. Tissues were put into 2 mL Eppendorf tubes with a pre-cooled 5 mm metal bead. The cryomill (Retsch, Newtown, PA) was pre-cooled before loading 2 mL tubes. All tissues were milled at 25 Hz for 1-2 minutes. Metal beads were removed, and tissue powder was weighed on dry ice.

### Metabolite extraction and measurements using LC-MS

For tissue metabolite extraction, cryomilled tissue powder (∼20 mg) was mixed with an extraction solvent (40:40:20 mixture of acetonitrile:methanol:water) to make 25 mg of tissue/mL of solvent, vortexed, and centrifuged at 16,000 x g for 10 min at 4°C. 3 μL of supernatant was injected to LC-MS. For serum, 150 μL of extraction solvent was added to 5 μL of serum and processed the same as above tissue extraction. A quadrupole orbitrap mass spectrometer (Q Exactive Plus; ThermoFisher Scientific) operating in negative or positive ion mode was coupled to a Vanquish UHPLC system (ThermoFisher Scientific) with electrospray ionization and used to scan from m/z 70 to 1,000 at 2 Hz, with a 140,000 resolution. LC separation was achieved on an XBridge BEH Amide column (2.1 x 150 mm^2^, 2.5 μm particle size, 130 Å pore size; Waters Corporation) using a gradient of solvent A (95:5 water: acetonitrile with 20 mM of ammonium acetate and 20 mM of ammonium hydroxide, pH 9.45) and solvent B (acetonitrile). Flow rate was 150 μl/min. The LC gradient was: 0 min, 85% B; 2 min, 85% B; 3 min, 80% B; 5 min, 80% B; 6 min, 75% B; 7 min, 75% B; 8 min, 70% B; 9 min, 70% B; 10 min, 50% B; 12 min, 50% B; 13 min, 25% B; 16 min, 25% B; 18 min, 0% B; 23 min, 0% B; 24 min, 85% B; and 30 min, 85% B. The autosampler temperature was 5°C and the injection volume was 3 μl. Metabolite concentrations were determined by authentic synthesized standards from Sigma by generating standard curves. Data were analyzed using the MAVEN software (build 682, http://maven.princeton.edu/index.php) and Compound Discoverer software (Thermofisher Scientific).

### Glycerol measurements using LC-MS

Enzymatic derivatization of glycerol to glycerol-3-phosphate was conducted using ATP (Abcam, AB146525) and glycerol kinase (Sigma-Aldrich, G6142-1KU), adapted from the previously published protocol^26^. Enzyme Mix was prepared by resuspending purified glycerol kinase from *Cellulomonas sp* in Derivatization Buffer (25 mM Tris-HCl pH 8, 50 mM NaCl, 10 mM MgCl_2_, 5 mM ATP, and 10% sucrose) to a concentration of 40 U/mL. 5 μL of serum was mixed with 2.5 μL Enzyme Mix and incubated at room temperature for 10 min. Enzymatic reactions were quenched by adding 950 μL 40:40:20 of acetonitrile: methanol: water + 0.1 M formic acid to each reaction and immediately neutralized with 88 μL of 15% NH_4_HCO_3_. Samples were vortexed and centrifuged at 16,000 x g for 10 min at 4°C. 3 μL of supernatant was injected for LC-MS analysis using the same setting for metabolomics above.

### RNA-sequencing analysis

RNA was isolated from kidney tissues using Qiazol (QIAGEN) and an RNeasy kit (QIAGEN) and RNA sequencing was performed at the UCI Genomics Core. Raw sequencing data was assessed for quality using FastQC and trimmed of adapter sequences using Trim Galore with the –paired setting. The trimmed paired end reads were aligned to the mm10 reference genome using STAR with the following settings: –outFilterMismatchNmax 2 -- outFilterMultimapNmax 4 --chimScoreSeparation 10 --chimScoreMin 20 --chimSegmentMin 15 --outSAMattributes All --outSAMtype BAM SortedByCoordinate. The resulting bam file was filtered for primary alignments using samtools with the flag 0X100. Multimappers were removed from the alignment file using samtools with setting -bq 255. Reads were quantified using HOMER’s analyzeRepeats with the following settings: -count exons -strand + -condenseGenes -norm 1e7.

### Statistics and Reproducibility

Heatmaps were generated using Metaboanalyst and R software (gplots). Statistical analysis was performed using Graphpad Prism 9.0 and R software (rstatix). When two groups were compared, a two-tailed, unpaired Student’s t-test was used to calculate P values, with P < 0.05 used to determine statistical significance. When >2 groups were compared, a one-way ANOVA was employed. Tukey’s method was used to correct for multiple comparisons. For correlation analysis, Pearson correlation test was used.

## Data and materials availability

All data needed to evaluate the conclusions in the paper are present in the paper. R scripts to calculate the carbon and nitrogen trafficking are available on GitHub: https://github.com/johnnl15/Bootstrapping_AUC_NitrogenFL_BATLiverSerum

## Acknowledgment

This paper is dedicated to Dr. Gina Lee, an incredibly kind and inspiring scientist and mentor who remains an important influence on our work. She will be missed dearly. We also thank all members of the Martinez, Jang and Lee laboratories for the discussion. Cartoon illustrations were created with BioRender.com.

## Author contributions

C.J. and G.L. conceived the project and supervised the study. M.E.K., L.A.H., and A.L.A performed sample processing and LC-MS analysis for the AV experiments. C.B.R. and J.K. performed AV blood sampling with the support from L.A.H and A.L.A. M.E.K and G.T. performed RNA sequencing analysis with the supervision by T.F.M. Y.K contributed to glycerol measurement and manuscript editing. W.C., K.J., Y.H.A., S.J., J.L., I.T., M.L.L., H.B., Y.C., and W.S.S., contributed to sample preparation, LC-MS maintenance, data analysis and intellectual discussion. M.E.K, C.J., and G.L. wrote the manuscript and all co-authors read and commented.

