## Supplementary Figures for "Sex-dependent metabolic remodeling of kidneys revealed by arteriovenous metabolomics"

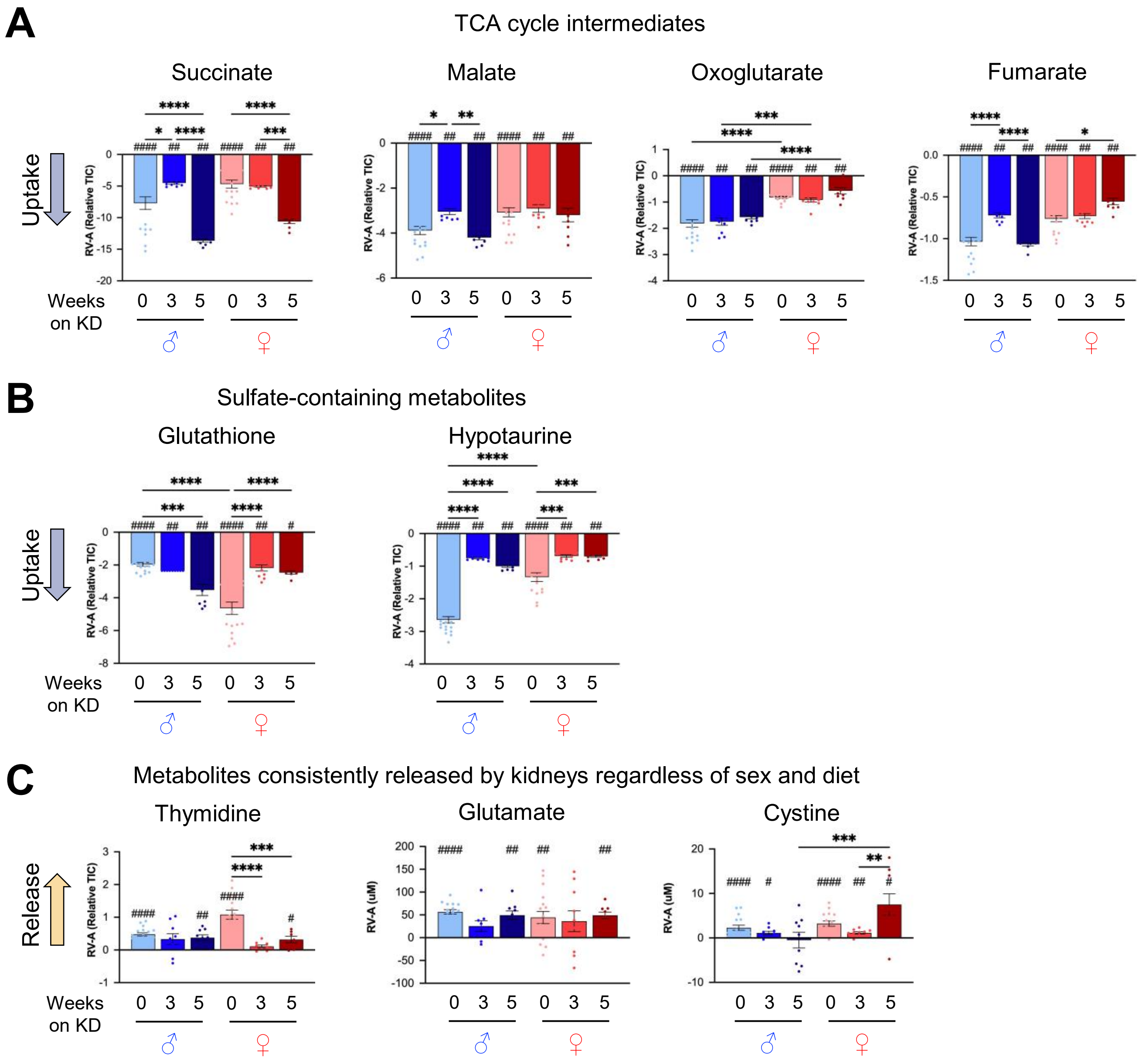


**Figure S1. Metabolite trafficking during diet switching in male and female kidneys.** (A-C) Kidney release and uptake of the indicated circulating metabolites. N=18, 9, and 9 mice for 0, 3 and 5 weeks KD-fed males and N=17, 10, and 9 mice for 0, 3, and 5 weeks KD-fed females. *$p<0.05$, **$p<0.01$, ***$p<0.001$, ****$p<0.0001$ by Students' t-test. $^{\#}p<0.05$, $^{\#\#}p<0.01$, $^{\#\#\#\#}p<0.0001$ by one-sample t-test.

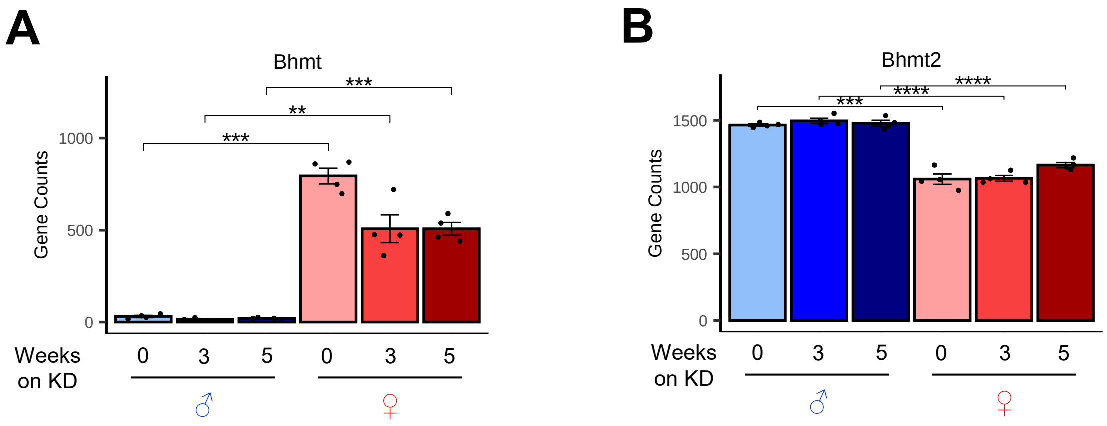


**Figure S2. Sex differences in gene expression of *Bhmt* and *Bhmt2*.** (A, B) RNA-seq gene counts of *Bhmt* or *Bhmt2* in male and female kidneys under NC or KD. N=4 mice per group. **$p<0.01$, ***$p<0.001$, ****$p<0.0001$ by one-way ANOVA.

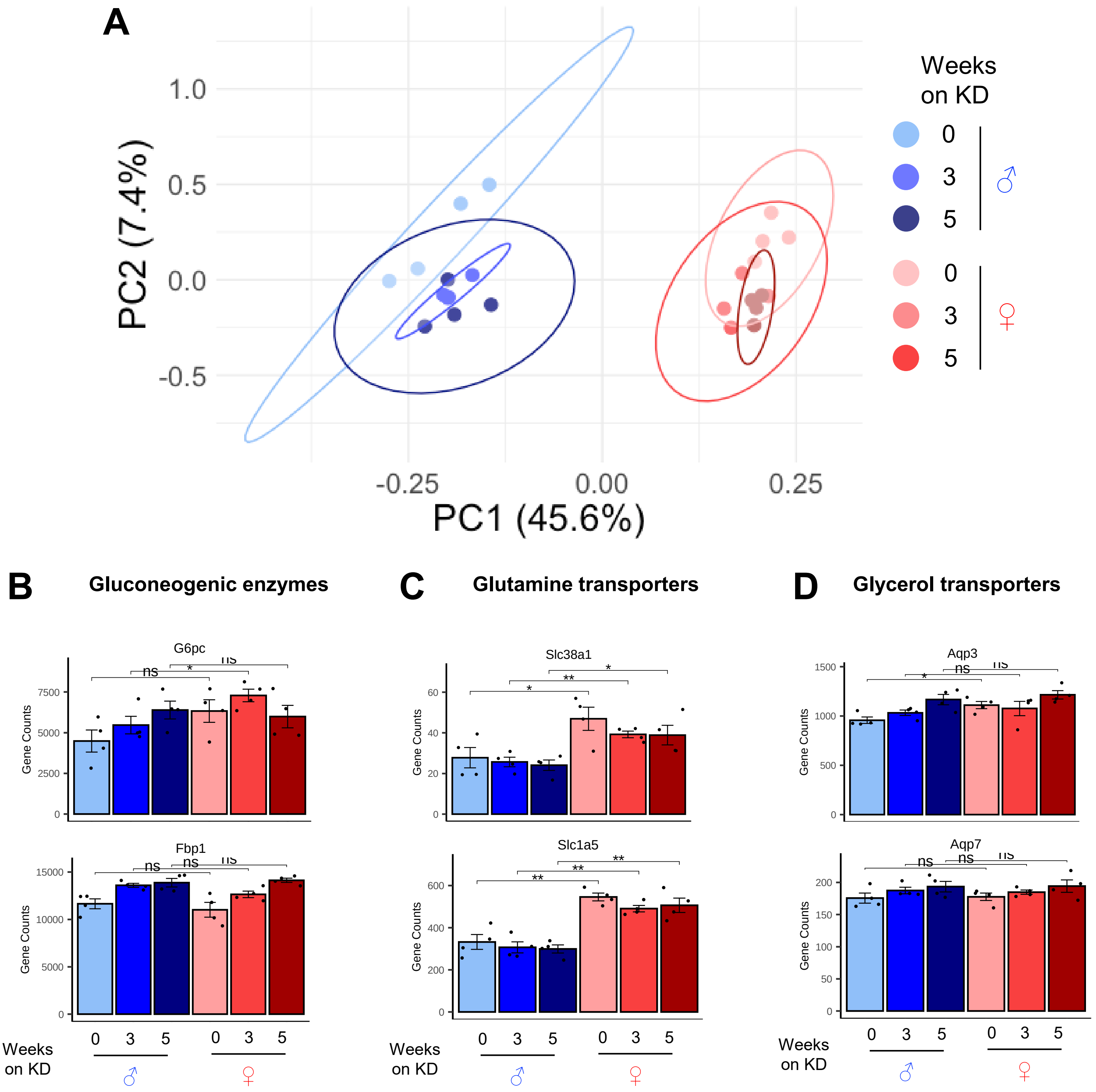


**Figure S3. Sex differences in kidney transcriptome under NC and KD.** (A) Principal component analysis (PCA) of RNA-seq results. (B-D) RNA-seq gene counts of the indicated genes. N=4 mice per group. *p<0.05, **p<0.01, by one-way ANOVA.
